# Dynamics of Fanconi anemia protein D2 in association with nuclear lipid droplet formation

**DOI:** 10.1101/2025.02.05.636583

**Authors:** Tomoya Hotani, Motonari Goto, Yukie Otsuki, Shun Matsuda, Nobuhiro Wada, Masakazu Shinohara, Tomonari Matsuda, Masayuki Yokoi, Kaoru Sugasawa, Yuki Ohsaki, Wataru Sakai

**Affiliations:** Biosignal Research Center, Kobe University, Kobe 657-8501, Japan; Department of Biology, Graduate School of Science, Kobe University, Kobe 657-8501, Japan; Research Center for Environmental Quality Management, Graduate School of Engineering, Kyoto University, Otsu 520-0811, Japan; Department of Environmental Engineering, Graduate School of Engineering, Kyoto University, Kyoto 615-8530, Japan; Department of Anatomy I, School of Medicine, Sapporo Medical University, Sapporo 060-8556, Japan; Division of Molecular Epidemiology, Kobe University Graduate School of Medicine, Kobe 650-0017, Japan; The Integrated Center for Mass Spectrometry, Kobe University Graduate School of Medicine, Kobe 650-0017, Japan

## Abstract

Fanconi anemia (FA) is a rare genetic disease caused by the loss of function of one of the 22 associated genes and is characterized by bone marrow failure, cancer predisposition, and developmental defects. The proteins encoded by these genes (FA proteins) mainly function in DNA damage response and repair. Although FA deficiency has multiple effects on the regulation of lipid metabolism, the molecular function of FA proteins in the context of FA pathology remains unclear. In the present study, we demonstrated that FANCD2, a key component of FA proteins, interacts with lipid metabolism-related factors and that FANCD2 deficiency downregulates the cellular levels of fatty acids. Moreover, a portion of FANCD2 is localized to nuclear lipid droplets in response to oleic acid treatment. These subcellular dynamics are independent of FANCD2 monoubiquitination, which is essential for the DNA damage response. Collectively, these findings demonstrate that FANCD2 responds to not only DNA damage but also oleic acid exposure, providing insights into the pathogenesis of lipid dysregulation in FA.

## Introduction

Fanconi anemia (FA) is a rare genetic disease in humans characterized by bone marrow failure, cancer predisposition, and developmental defects (Auerbach, 2009). To date, more than 20 genes have been identified to be responsible for FA (Niraj et al., 2019). The products of these genes (FA proteins) are mainly involved in the maintenance of genome stability, particularly in DNA interstrand crosslink (ICL) repair, which is known as the FA pathway. Nine FA proteins and other FA-related factors form an FA core complex with E3 ubiquitin ligase activity, which functions as an upstream regulator of the FA pathway. In response to DNA damage, FANCD2 and FANCI form a heterodimer—known as the ID complex—and are monoubiquitinated by the FA core complex. This reaction is a crucial step in orchestrating the entire FA pathway and the downstream repair process. Over recent decades, FA proteins and related factors have been isolated, revealing the molecular mechanisms underlying the FA pathway. Recent studies have shown that endogenous aldehydes have a significant effect on FA symptoms (Garaycoechea et al., 2012; Langevin et al., 2011; Pontel et al., 2015). Defects in FA proteins result in the dysregulation of aldehyde-induced DNA damage response and repair, eventually leading to FA symptoms such as bone marrow failure and cancer predisposition. Moreover, endocrine and metabolic abnormalities are reported as one of the common phenotypes of FA (Giri et al., 2007; Petryk et al., 2015). Approximately 80% of patients with FA have at least one endocrine abnormality. More than half of the patients with FA have impaired lipid metabolism. Consistent with these FA symptoms, alterations in lipid metabolism resulting from FA deficiency have been reported at the cellular level. Mesenchymal stromal cells from *Fanca* and *Fancd2* knockout mice exhibited abnormal lipid profiles, indicating disturbances in fatty acid and lipid metabolism (Amarachintha et al., 2015), and *Fancd2*-deficient male mice exhibited altered hepatic lipid and bile acid metabolism when fed the Paigen diet (Moore et al., 2019). FA deficiency has been reported to affect glycosphingolipid metabolism (Zhao et al., 2018). Additionally, FA-deficient human cells show accumulation of lipid droplets (LDs) (Ravera et al., 2019), which constitute an organelle crucial for lipid metabolism. LDs are composed of lipid monolayers in which neutral lipids, triacylglycerol, and cholesterol ester are stored. Various factors related to lipid metabolism have been identified in the LD membrane (Bersuker et al., 2018), which play an important role in lipid metabolism, such as energy production, membrane trafficking between organelles, and lipid accumulation (Olzmann and Carvalho, 2019). Although LDs are classically defined as cytoplasmic organelles, it has recently been reported that LD formation also occurs in the nucleus in various cell types and tissues (Uzbekov and Roingeard, 2013; Wang et al., 2013; Layerenza et al., 2013; Ohsaki et al., 2016; Zadoorian et al., 2023; McPhee et al., 2024; Romanauska and Köhler, 2018, 2021), suggesting that lipid metabolism via nuclear LDs (nLDs) occurs even in the nucleus.

Increasing evidence suggests a functional link between FA proteins and lipid metabolism. However, the relevant molecular mechanisms underlying the link between FA deficiency and abnormal lipid metabolism remain unclear. In the present study, gene ontology analysis following screening for FANCD2-binding factors indicated that FANCD2 interacts with factors related to fatty acid and lipid metabolism. Furthermore, we found that FANCD2 localizes around nLDs in response to oleic acid exposure. These results suggest that FANCD2 responds not only to DNA damage but also to nLD formation.

## Results and discussion

### Lipid metabolism-related factors as FANCD2-interacting proteins

To obtain novel insights into FANCD2 function independent of the DNA damage response, we performed mass spectrometry using the human osteosarcoma U2OS cell line stably expressing FLAG epitope-tagged FANCD2 (FLAG-FANCD2) (Fig S1A). The cell lysate for mass spectrometry analysis was prepared from the cell culture without treatment with an exogenous DNA-damaging agent. The protein complex with FLAG-FANCD2 was subjected to SDS-PAGE followed by CBB staining (Fig S1B) and mass spectrometric analysis as described in the Materials and Methods. We detected specific bands from the FLAG-FANCD2 lysate compared with that of the control. In total, we identified 135 proteins as candidate FANCD2-interacting proteins (Supplementary Table 1). Previously, numerous proteins have been isolated as FANCD2-binding factors, many of which were screened under DNA damage conditions using DNA-damaging agents (Smogorzewska et al., 2007; Liu et al., 2010; Unno et al., 2014). However, the candidate proteins identified in our screening did not contain previously reported FANCD2-interacting proteins, including FANCI. This finding may be attributed to the lack of exposure to exogenous genotoxic stress in our screening. Consistent with this finding, FANCD2 and FANCI have been reported to form a weak complex under standard immunoprecipitation conditions (Sareen et al., 2012).

To explore the potential biological processes associated with the candidate proteins, gene ontology (GO) enrichment analysis was performed using DAVID Bioinformatics Resources (Sherman et al., 2022). Remarkably, the top-ranked annotation cluster was associated with fatty acid and lipid metabolism or biosynthesis (Fig 1A). There is increasing evidence for a link between FA deficiency and changes in lipid homeostasis (Degan et al., 2019). Notably, more than 50% of patients with FA are diagnosed with dyslipidemia associated with glucose intolerance or hyperglycemia (Petryk et al., 2015). To elucidate the direct role of FANCD2 in cellular lipid metabolism, a *FANCD2* knockout (FANCD2^KO^) cell line was generated from U2OS cells (Fig S1C), and the total fatty acid content in the cells was measured. As shown in Figure 1B, the total fatty acid content in FANCD2^KO^ cells was significantly lower than that in the parental U2OS cell line and was partially restored by the ectopic expression of wild-type FANCD2. The levels of stearic acid (18:0) among saturated fatty acids and 11-eicosenoic acid (20:1n-9), erucic acid (22:1n-9), and arachidonic acid (20:4n-6) among unsaturated fatty acids showed a significant decrease in FANCD2^KO^ cells and restoration in the complemented cells, respectively. The other fatty acids tested in the present study showed similar patterns, but the differences were not statistically significant.

**Figure 1.**
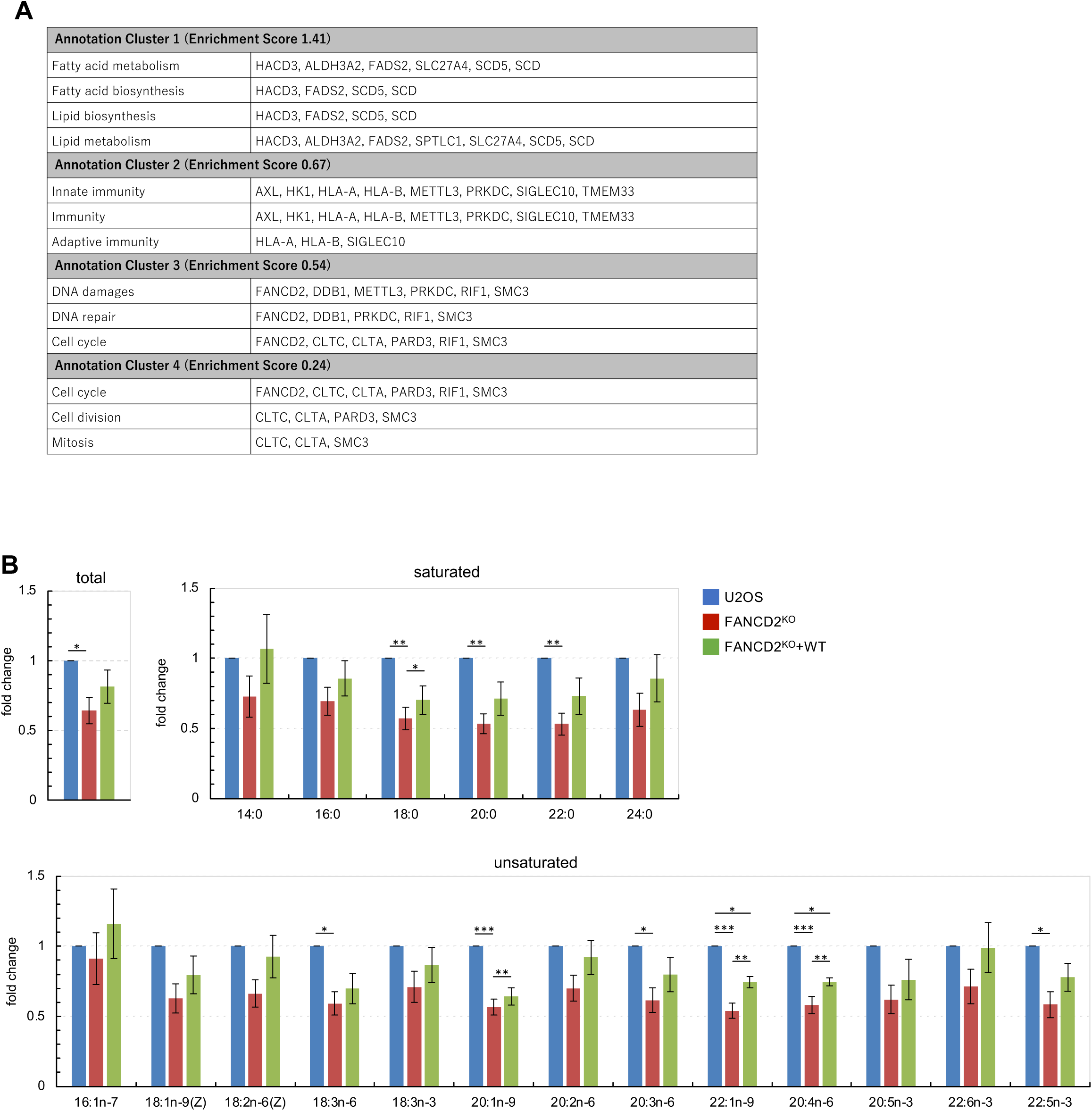
Evaluation of cellular fatty acid levels in FANCD2^KO^ cells. (**A**) Functional annotation clustering by biological process. The enrichment score represents the geometric mean (in −log_10_ scale) of the *p*-value in the corresponding annotation cluster. In this GO analysis, an enrichment score of 1.41 indicates a statistically significant difference (*p* < 0.05). (**B**) Cellular fatty acid levels were analyzed using gas chromatography/mass spectrometry (GC/MS). The relative levels of each fatty acid (saturated and unsaturated) and total fatty acids are shown (mean ± standard error of the mean (SEM) from 4 independent experiments, one-way analysis of variance coupled with Tukey’s multiple comparisons test, \**p* < 0.05, \*\**p* < 0.01, \*\*\**p* < 0.001). Blue: U2OS cells, red: FANCD2^KO^ cells, green: complemented FANCD2^KO^ cells (FANCD2^KO^+WT). Supplementary Table 2 presents the absolute data.

### Dynamics of FANCD2 in response to oleic acid exposure

The molecular mechanism underlying the association between FANCD2 and fatty acid metabolism remains poorly understood. To test the hypothesis that FANCD2 functions in not only the DNA damage response but also fatty acid metabolism, we analyzed the subcellular localization of EGFP-FANCD2 (Fig S1D) under conditions of increased fatty acid levels. In a culture medium supplemented with oleic acid (OA), most of the FANCD2 retained its localization in the nucleus, as observed under normal culture conditions. However, live-cell imaging captured the atypical nuclear dynamics of FANCD2 after OA treatment. Confocal microscopy showed that a portion of FANCD2 exhibited “ring-shaped” localization. In some cells, the ring-shaped FANCD2 grew gradually from small dots (Fig 2A). In another cell, the ring-shaped FANCD2 gradually disappeared (Fig 2B). The dynamics of the ring-shaped FANCD2 were indicative of nuclear lipid droplet (nLD) formation upon OA treatment. Under OA culture conditions, nLD formation occurs rapidly in U2OS cells (Ohsaki et al., 2016; Sołtysik et al., 2021; Lee et al., 2020). Double fluorescence labeling revealed unique subcellular localization of FANCD2 around nLDs in the OA-supplemented medium (Fig 2C). This LD localization of FANCD2 was enhanced in a dose-dependent manner (Fig 2D), and its maximal induction was detected after 2 days of OA treatment (Fig 2E). Moreover, some LDs with FANCD2 were detected even in the cytoplasm (Fig S2A). It is difficult to distinguish whether these LDs with FANCD2 were observed as a result of *de novo* localization of FANCD2 in cytoplasmic LDs or sustained localization of FANCD2 in nLDs beyond cell division. It has previously been reported that nLDs are extruded into the cytoplasm after mitosis (Sołtysik et al., 2019); thus, FANCD2 may also maintain its localization in nLDs. To examine the nLD localization of endogenous FANCD2, immunofluorescence analysis was performed using anti-FANCD2 antibody. The results of this analysis also demonstrated the nLD localization of endogenous FANCD2 in the OA-supplemented medium (Fig 2F). Furthermore, correlative light–electron microscopy (CLEM) showed that LDs containing FANCD2 obviously existed in the nucleoplasm and were not cytoplasmic LDs inside the invagination of the nuclear membrane (Fig 2G,H). No membranous structures were observed on the surface of LDs with FANCD2. Notably, some LDs with FANCD2 interacted with nucleoli or condensed chromatin structures (Fig 2H iv and Fig S2C iii), whereas others did not (Fig 2H iii and Fig S2C ii). We and others previously found that nLDs are often associated with PML proteins and PML body components (Ohsaki et al., 2016; Lee et al., 2020). PML bodies are subnuclear domains that interact with chromatin and regulate various cellular responses such as gene expression (Kurihara et al., 2020). Moreover, nLDs are reported to be involved in the transcriptional regulation in yeast cells and mammalian cells (Romanauska and Köhler, 2018; Umaru et al., 2023). These findings prompted us to verify the nuclear localization of FANCD2 and PML following OA treatment. Double-labeled immunofluorescence staining showed that FANCD2 and PML coexisted on some nLDs after OA treatment (Fig S2D). However, further analysis is required to determine whether FANCD2 localization affects the association of nLDs with chromatin and subsequently modify gene expression.

**Figure 2.**
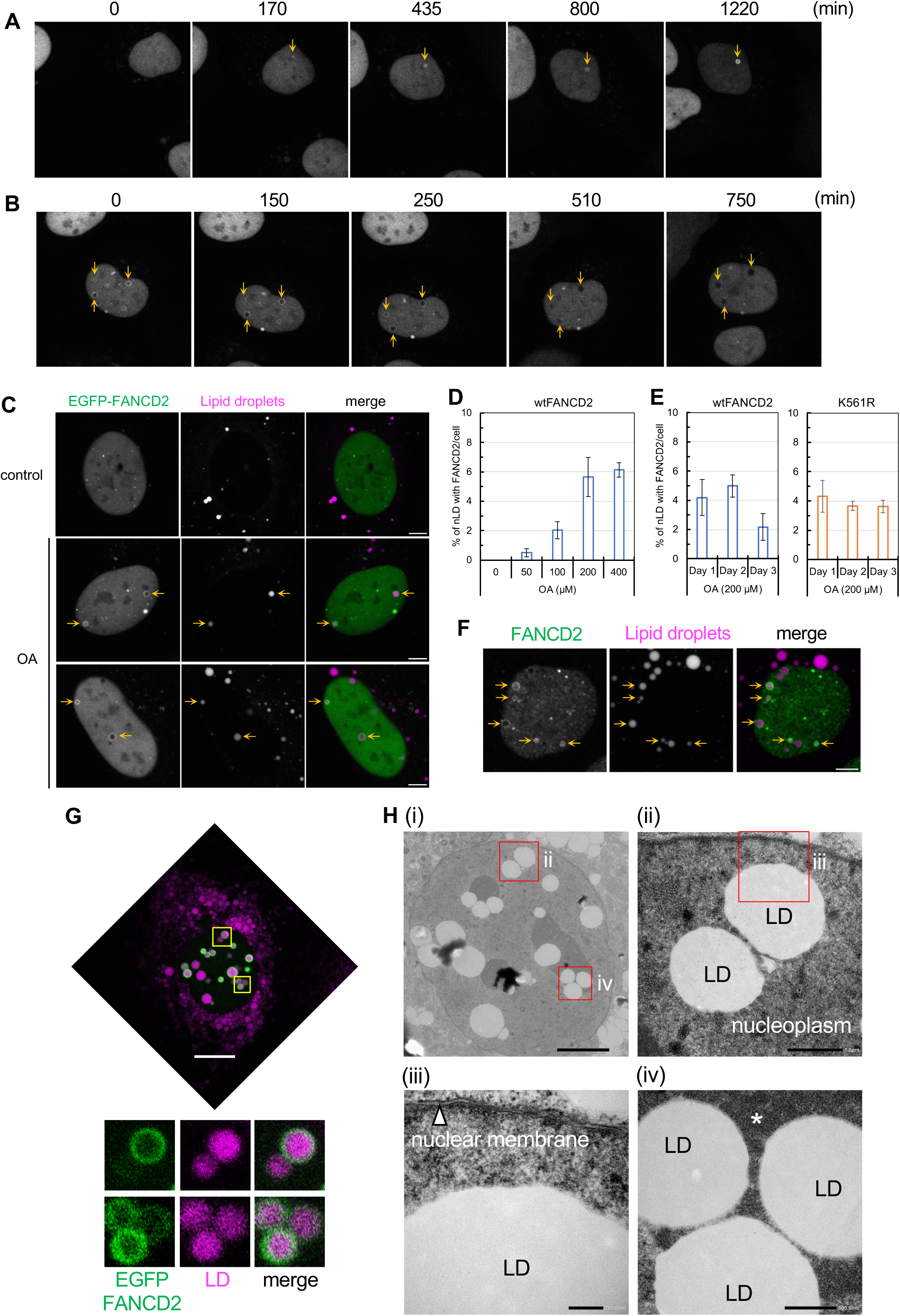
FANCD2 localizes around nuclear lipid droplets. **(A, B)** Images of EGFP-FANCD2 in medium supplemented with 200 µM oleic acid (OA) are shown. **(A)** Growth of EGFP-FANCD2 from small dots to a ring-like appearance (yellow arrows). Selected frames from Video 1 are shown. **(B)** Disappearance of ring-like EGFP-FANCD2. Selected frames from Video 2 are shown. **(C)** Localization of EGFP-FANCD2 in cells cultured with 200 µM of OA for 2 days. LDs were stained with Lipi-Blue. Scale bars, 5 µm. **(D, E)** Bar graph showing the percentage of nLDs with FANCD2 in the cell (mean ± SEM from more than 3 independent experiments). **(F)** Localization of endogenous FANCD2 in U2OS cells cultured with 200 µM of OA for 2 days. Scale bar, 5 µm. **(G, H)** Correlative light–electron microscopy. U2OS cells stably expressing EGFP-FANCD2 were treated with 100 µM of OA for 24 h. **(G)** Cells were weakly fixed, and fluorescence images were captured first. Some of the nLDs (magenta) were surrounded by FANCD2 (green). Highly magnified images (yellow squares) are shown below. The upper and lower panels correspond to the images in H (ii) and (iv), respectively. Scale bar, 10 µm. **(H)** (i) Electron microscopy images of the cell shown in **(G)**. (ii) (iii) (iv) Highly magnified images (red squares) are shown. Scale bars, 5 µm (i), 1 µm (ii), 200 nm (iii), and 500 nm (iv). Asterisk indicates the nucleolar area.

### Oleic acid-induced nuclear lipid droplet formation does not stimulate the DNA damage response

To examine whether the induction of nLD formation with OA leads to a DNA damage response, the phosphorylation of histone H2AX at serine 139 (γH2AX), a molecular marker of DNA damage, was detected by immunoblot analysis. As shown in Fig 3A, OA treatment had no effect on the level of γH2AX, unlike treatment with the DNA crosslinker mitomycin C (MMC). Consistent with this finding, FANCD2 monoubiquitination was not induced under the indicated culture conditions (Fig 3B). In line with previous reports (Urahama et al., 2008; Liu et al., 2019), cell viability under OA treatment (100–400 µM) was not significantly different from that under normal culture conditions (data not shown). Furthermore, immunofluorescence analysis using an anti-γH2AX antibody showed that γH2AX signals did not colocalize with nLD and FANCD2 in the OA-supplemented medium (Fig 3C, middle panels). In contrast, MMC treatment had no effect on LD formation, but most nuclear foci of γH2AX colocalized with FANCD2 (Fig 3C, bottom panels), indicating the sites of MMC-induced DNA damage. The FANCD2 ubiquitin-dead K561R mutant (Fig S1D), which can no longer be monoubiquitinated by the FA core complex (Garcia-Higuera et al., 2001), still exhibited nLD localization in the OA-supplemented medium, similar to the wild-type FANCD2 (Fig 2E). These findings indicate that the nLD localization of FANCD2 is independent of the canonical DNA damage response of FANCD2.

**Figure 3.**
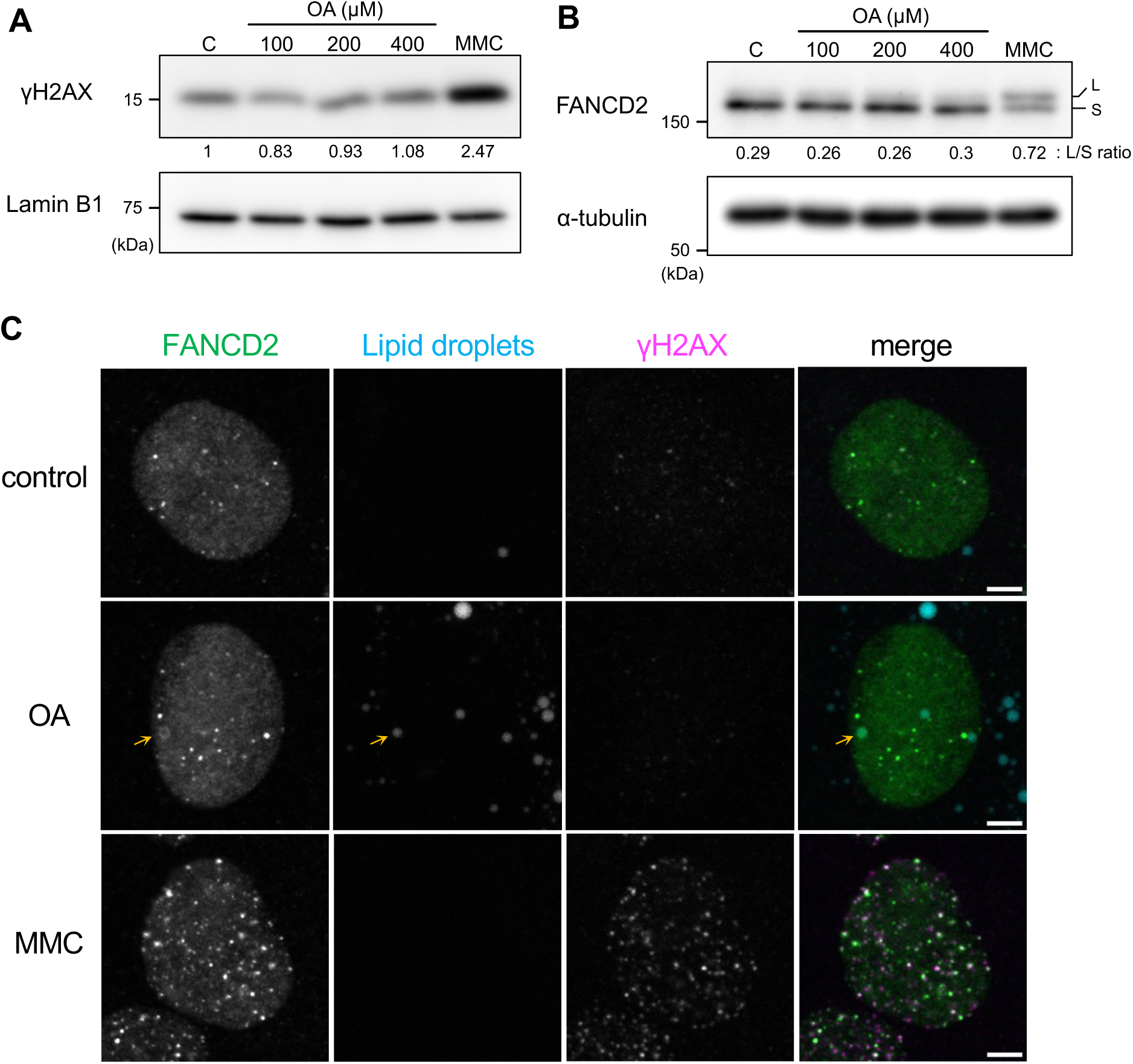
Nuclear lipid droplet localization of FANCD2 is independent of DNA damage response. **(A)** U2OS cells were cultured with the indicated concentration of oleic acid (OA) for 2 days. DNA damage was induced by treating cells with 500 nM mitomycin C (MMC) for 24 h. The fold change in γH2AX levels relative to the control is shown. Lamin B1 was used as a loading control. **(B)** The conditions for OA and MMC treatment were the same as those described in **(A)**. Values below the FANCD2 panel depict the L/S ratios between the monoubiquitinated form (L) and unmodified form (S) of FANCD2. α-tubulin was used as a loading control. **(C)** Immunofluorescence analysis of FANCD2 (green) and γH2AX (magenta). LDs (cyan) were stained with Lipi-Blue. Cells were exposed to MMC (150 nM) overnight or OA (200 µM) for 2 days. Orange arrows indicate an nLD with FANCD2. Representative images are shown. Scale bar, 5 µm.

### Oleic acid-induced nuclear lipid droplet localization of FANCI

In response to DNA damage, monoubiquitinated FANCD2 and FANCI form a stable heterodimeric complex (ID complex) that targets the site of DNA damage. To examine whether FANCI and FANCD2 colocalize around nLDs after OA treatment, we generated cells stably expressing each protein labeled with different fluorescent proteins, mCherry and EGFP, respectively (Fig S1E). Consistent with previous findings (Smogorzewska et al., 2007), in response to MMC exposure, FANCD2 formed nuclear foci, which were consistent with those of FANCI (Fig 4A, bottom panels). However, unlike FANCD2, FANCI exhibited no nLD localization under this condition (Fig 4A, middle panels). However, nLD localization of FANCI was detected after FANCD2 depletion in the same cells (Fig 4B–D). These results indicate that both FANCD2 and FANCI can localize to nLDs, but FANCD2 has a higher tendency to localize to nLDs. The findings suggest that, unlike during the DNA damage response, the two proteins do not stably interact on nLDs to form an ID complex. Consistent with this notion, OA treatment did not promote the complex formation of FANCD2 with FANCI (Fig S3A), unlike MMC treatment (Fig S3B). Alternatively, it is likely that their localization to nLDs is competitive (see below).

**Figure 4.**
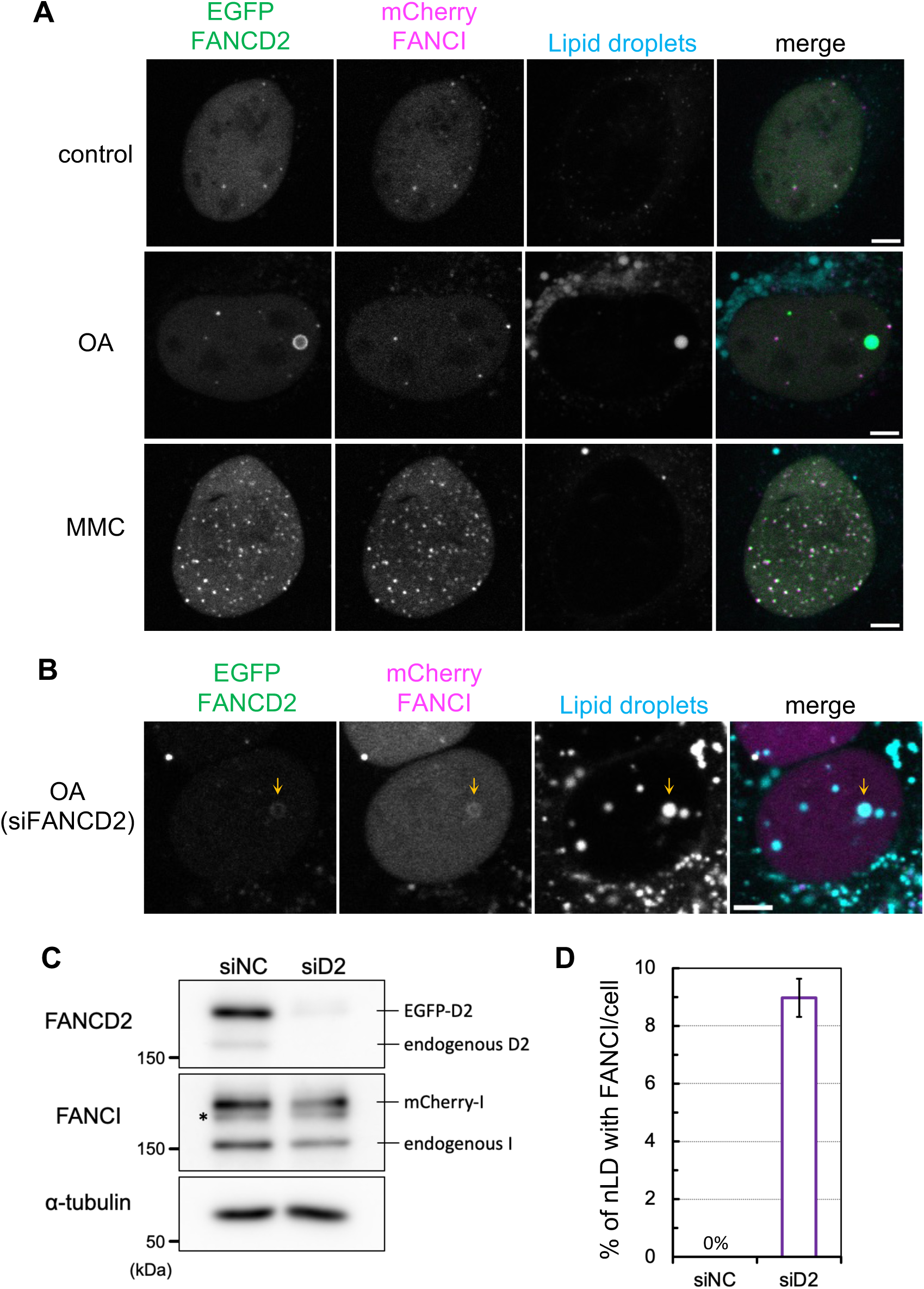
Oleic acid-induced nuclear lipid droplet localization of FANCI. **(A)** Live-cell imaging of U2OS cells expressing both EGFP-FANCD2 and mCherry-FANCI. Lipid droplets (LDs) were stained with Lipi-Blue. Representative images are shown. Scale bar, 5 µm. The conditions for oleic acid (OA) and mitomycin C (MMC) treatment were the same as those described in Figure 3C. **(B)** LD localization of FANCI after FANCD2 depletion. The culture conditions were the same as those described in Figure 2C. Orange arrows indicate FANCD2 and FANCI colocalization in the nLD. Scale bar, 5 µm. **(C)** Immunoblot analyses confirming the depletion of FANCD2 proteins. Asterisk indicates monoubiquitinated endogenous FANCI. **(D)** Bar graph showing the percentage of nLDs with FANCI (mean ± SEM from more than 3 independent experiments). siNC, negative control siRNA; siD2, siFANCD2.

### Mobility of FANCD2 on nuclear lipid droplets

Two types of subnuclear localization of FANCD2 have been identified: DNA-damage-induced nuclear foci formation and OA-induced nLD localization. However, the stability of both types remains unclear. To investigate the mobility of the nuclear localization of FANCD2, fluorescence recovery after photobleaching (FRAP) analysis was performed using cells stably expressing EGFP-FANCD2 (Fig 5). Nucleoplasmic FANCD2 (no subnuclear localization) showed rapid recovery kinetics under both culture conditions, whereas FANCD2 on nLDs and DNA-damage-induced FANCD2 foci showed significantly lower recovery kinetics. Collectively, these findings demonstrate that FANCD2 undergoes dynamic changes in subnuclear localization in response to OA treatment. Nevertheless, the mobility of FANCD2 around nLDs was limited, indicating a tendency toward stable localization.

**Figure 5.**
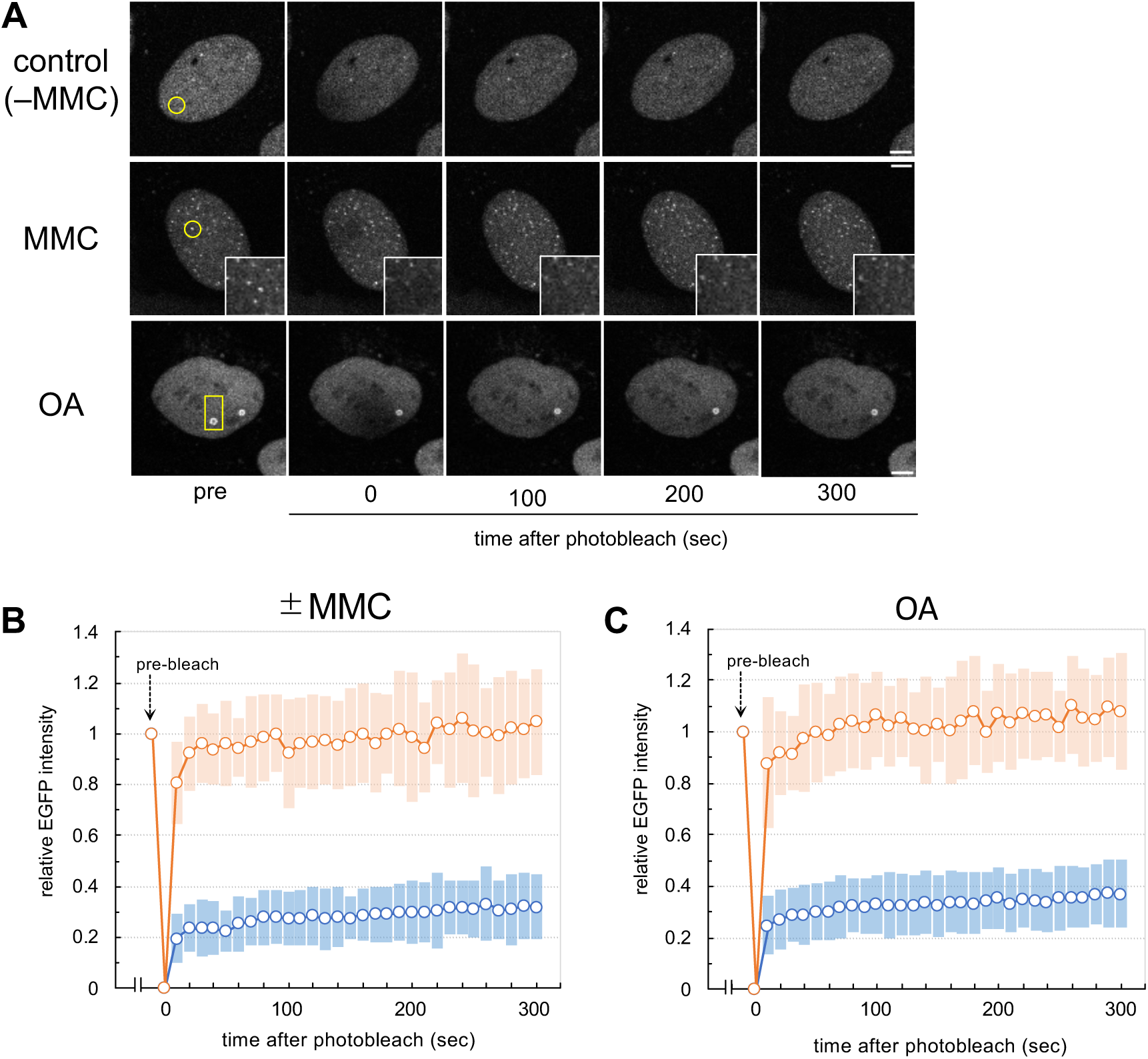
Mobility of FANCD2 on nuclear lipid droplets. **(A)** Selected frames are shown. The yellow circle and rectangle indicate the photobleach area. The EGFP intensity in each bleached area was calculated and plotted over time (mean ± standard deviation; see Fig S2 E,F). The highly magnified images in the middle panels (MMC) represent the bleached areas of the FANCD2 foci. The conditions for oleic acid (OA) and mitomycin C (MMC) treatment were the same as those described in Figure 3C. Scale bars, 5 µm. **(B)** Fluorescence recovery after photobleaching (FRAP) analysis of EGFP-FANCD2 expressed in U2OS cells (±MMC). Blue and orange symbols indicate the fluorescence recovery of FANCD2 nuclear foci (+MMC) and nucleoplasmic FANCD2 under normal culture conditions (-MMC), respectively. **(C)** FRAP analyses of EGFP-FANCD2 expressed in U2OS cells (+OA). Blue and orange symbols indicate the fluorescence recovery of FANCD2 in nLDs and nucleoplasmic FANCD2, respectively.

Over the past 30 years, the various molecular functions of FA proteins have been identified. FA proteins play an important role in the maintenance of genomic stability, particularly in ICL damage response and repair. However, although more than half of patients with FA exhibit impaired endocrine regulation and lipid metabolism, the mechanism by which FA protein deficiency causes dysregulation remains unclear. Although several studies have shown that the FANCD2 protein interacts with various factors in response to DNA damage, the present findings indicate that FANCD2 interacts with lipid-metabolism–related factors under conditions without exogenous DNA damage (Fig 1A). FANCD2^KO^ cells showed downregulation of cellular fatty acid levels in normal culture conditions (Fig 1B). Previously, lipidomics profiling of FA-deficient cells indicated altered lipid metabolism, particularly a reduction in neutral lipid synthesis and the accumulation of sphingolipids such as gangliosides and sphingomyelins (Zhao et al., 2018; Moore et al., 2019). In the present study, metabolic profiling was used to detect cellular fatty acids from free fatty acids and neutral lipids that form ester bonds. However, amide-bound fatty acids in sphingolipids were not targeted (see Material and Methods). In FA-deficient cells, intracellular fatty acids may be used for sphingolipid synthesis, resulting in reduced total fatty acid content in neutral lipids. Notably, our GO analysis revealed that the top-ranked annotation cluster was related to lipid and fatty acid metabolism (Fig 1A). HACD3, FADS2, SCD5, and SCD are involved in fatty acid biosynthesis, whereas SPTLC1 and ALDH3A2 function in the *de novo* synthesis and degradation pathways of sphingolipid metabolism, respectively. These results suggest that FANCD2 is involved in the regulation of fatty acid biosynthesis and/or sphingolipid metabolism.

Moreover, a portion of FANCD2 localized around nLDs in the OA-supplemented culture medium (Fig 2). The ubiquitination of FANCD2 was not induced under these conditions (Fig 3). Consistent with this finding, the ubiquitin-dead K561R mutant exhibited nLD localization (Fig 2E), suggesting that FANCD2 localization on nLDs is independent of the canonical FA DNA damage response and repair pathway. In line with this notion, the colocalization of FANCD2 and FANCI was not essential for nLD localization (Fig 4). These are novel cellular responses of FANCD2 upon exposure to OA independent of the DNA damage response. The biological and physiological significance of FANCD2 in nLDs require further exploration. The canonical cellular location for lipid metabolism is cytoplasmic organelles such as the endoplasmic reticulum, mitochondria, and cytoplasmic LDs. These organelles communicate via local membrane contacts to promote lipid synthesis and lipolysis. Intriguingly, recent studies have reported that lipid metabolism occurs in the inner nuclear membrane (INM) and nLDs of both mammalian and yeast cells (Romanauska and Köhler, 2018; Lee et al., 2020; Sołtysik et al., 2021). In yeast cells, nLD recruits Opi1—a transcription suppressor—and regulates lipid metabolism, including phosphatidylinositol lipids (Romanauska and Köhler, 2018). In mammalian cells, some nLDs are enriched in CTP:phosphocholine cytidylyltransferase alpha (CCTα, a rate-limiting enzyme of *de novo* phosphatidylcholine synthesis pathway) and LPIN1 (a phosphatidic acid phosphatase catalyzing the conversion from phosphatidic acid to diacylglycerol) (Sołtysik et al., 2019, 2021; Lee et al., 2020). Moreover, enzymes involved in lipid metabolism are distributed in the INM and nLDs (Prasad et al., 2011; Vieyres et al., 2020). The present study revealed the stable and dynamic properties of FANCD2 in nLDs, suggesting the novel possibility that FANCD2 regulates fatty acid and/or lipid metabolism in the nucleus. Alternatively, considering the role of FA proteins in cytoplasmic function in selective autophagy for organellar quality control (Sumpter et al., 2016), nLDs with FANCD2 may be responsible for organellar quality control such as lipophagy.

Nevertheless, this study has three limitations that can be addressed in future studies. First, this study focused on the cellular dynamics of FANCD2 following OA exposure. However, it remains unknown whether other FA proteins localize around nLDs, similar to FANCD2. Second, although FANCD2 localizes around nLDs upon OA exposure, all nLDs in the same nucleus do not show the presence of FANCD2, and a portion of FANCD2 dissociates from nLDs. This discrepancy may be explained as follows: (i) The interaction of FANCD2 with the nLD surface may be accelerated via unknown partner proteins or suppressed by a competitor such as FANCI. Previously, we and other researchers have shown that the recruitment of CCTα to nLDs is competitively regulated by PLIN3 (Sołtysik et al., 2019) or PML (Lee et al., 2020). (ii) Alternatively, the lipid composition on the surface of nLDs may affect the localization of FANCD2. For instance, the amount of phosphatidic acids on the surface of nLDs appeared to vary in minutes (Sołtysik et al., 2021), suggesting that FANCD2 is recruited to nLDs through fluctuating concentrations of lipid-binding partners. (iii) Another likely explanation is that the size of nLDs defines the association of FANCD2 with nLDs. Smaller nLDs tend to have “packing defects” in their phospholipid monolayer because of their higher curvature, allowing the proteins to interact with the hydrophobic region of nLDs, and then the proteins may be released as the nLDs grow in size. Third, it remains unknown whether the presence of FANCD2 on nLDs affects the function of LDs in lipid metabolism. FANCD2 on nLDs may directly regulate the activity of lipid metabolism enzymes identified as candidate FANCD2-binding partners in this study. Alternatively, these proteins may regulate the functions of transcriptional factors around nLDs, as previously reported (Romanauska and Köhler, 2018; Umaru et al., 2023). To overcome these limitations, the molecular mechanisms of FA proteins in lipid metabolism must be investigated in detail. Further research on FANCD2 and other FA proteins in nLDs will elucidate one of the fundamental questions in the pathophysiology of FA.

## Materials and methods

### Cell lines and cell culture

The human osteosarcoma cell line U2OS and its derivatives were cultured at 37°C in a humidified atmosphere containing 5% CO_2_ in Dulbecco’s modified Eagle medium (Shimadzu Diagnostics) supplemented with 10% fetal bovine serum. The retroviral expression vector pMMPpuro was used for the stable expression of FLAG-FANCD2, EGFP-FANCD2, and mCherry-FANCI. The pMMPpuro empty vector was used to establish the control cell line (+vec). The production of recombinant retroviruses and infection were performed as previously described (Sakai and Sugasawa, 2014). A culture medium containing 1 µg/ml puromycin (Merck) was used to select stable transformants. Mitomycin C was purchased from Fujifilm Wako (139-18711).

### Preparation of the FANCD2 complex

All procedures were performed at 4°C or on ice. To prepare the FANCD2 complex, cell extracts were prepared with CSK buffer (10 mM PIPES-NaOH [pH 6.8], 3 mM MgCl_2_, 1 mM EGTA, 150 mM NaCl, 10% glycerol, 0.1% Triton X-100, 50 mM NaF) containing protease inhibitors (0.25 mM phenylmethylsulfonyl fluoride, 1 µg/ml leupeptin, 2 µg/ml aprotinin, 1 µg/ml pepstatin, 50 µg/ml Pefabloc SC). After incubation for 60 min, the cell lysates were centrifuged for 10 min at 20,000 × *g* to obtain soluble extracts. The FANCD2 complex was immunoprecipitated from the soluble extracts by incubating with anti-DYKDDDDK tag antibody beads (Fujifilm Wako) overnight with rotation. After extensive washes with CSK buffer, the bound proteins were eluted from the beads via incubation for 60 min with 0.5 mg/ml FLAG peptide (Fujifilm Wako) in the same buffer. After ammonium sulfate precipitation, the precipitated proteins were dissolved in SDS sample buffer (62.5 mM Tris-HCl [pH 6.8], 10% glycerol, 1% SDS, 1% 2-mercaptoethanol) and dialyzed with Biotech CE MWCO 8000 (Bio-tech) in the same buffer. Samples were separated by SDS-PAGE, and the CBB-stained bands were excised and digested in-gel with trypsin.

### Nano-liquid chromatography–mass spectrometry (LC-MS) and database search

The desalted samples were then subjected to LC-MS analysis using a Q-Tof Ultima API (Waters) coupled to a NanoFrontier nLC (Hitachi). The peptides were resolved by reversed-phase chromatography using a homemade capillary column (10 cm long, 50 μm in diameter) packed with 3 µm silica, using ReproSil-Pur C18-AQ resin (Dr Maisch GmbH). Peptides were eluted as follows (solvent A, 2% acetonitrile and 0.1% formic acid; solvent B, 98% acetonitrile and 0.1% formic acid; flow rates: 200 nl/min): 0–5 min, linear gradient from 2% to 5% B; 5–60 min, linear gradient to 23% B; 60–65 min, linear gradient to 70% B; 65–70 min, isocratic with 70% B; 75–75.1 min, linear gradient to 2% B; 75.1–105 min, isocratic with 2% B. MS was performed in positive mode, and spectra were acquired from 300 to 1800 m/z for 1 s followed by three data-dependent MS/MS scans from 45 to 900 m/z for 2 s each. The collision energy used to perform MS/MS was automatically varied according to the mass and charge state of the eluting peptide. Only the spectra of ions with charge states 2, 3, and 4 were acquired for the MS/MS analysis. The dynamic exclusion duration was set to 60 s. The typical mass spectrometric conditions were as follows: capillary voltage, 3.2 kV; source temperature, 80°C. Matrix Science Mascot Distiller and Mascot Server were used for peptide peak picking and protein identification, respectively. Database search was performed against the human International Protein Index protein database version 3.53) with the following parameters: enzyme, trypsin; missed cleavages, 2; fixed modification, carbamidomethyl (C), variable modification, methionine oxidation, pyroglutamylation (N-terminal E); peptide length, more than 6. A peptide mass tolerance of 100 ppm and fragment mass tolerance of 0.3 Da were used. The false discovery rate was set to <0.05. Gene ontology analysis was performed using DAVID Bioinformatics Resources (Sherman et al., 2022).

### Gene disruption of *FANCD2*

Disruption of the endogenous *FANCD2* gene in the U2OS cell line was performed using the GeneArt CRISPR Nuclease Vector with CD4 Enrichment Kit (Thermo Fisher Scientific). The gRNAs targeted sequences within exon 11 of FANCD2 (5’-GTTGTCGTCTATTAGATTGG-3’). After enrichment with anti-CD4 magnetic beads, single clones were isolated by limiting dilution. FANCD2 expression in each clone was assessed using immunoblotting, and gene disruption was confirmed using Sanger sequencing of genomic DNA.

### Total fatty acid analysis with gas chromatography/mass spectrometry (GC/MS)

For GC/MS analysis, cells were collected with 70% methanol after two rinses with PBS. The protein concentration in each cell extract was determined and normalized. Nonadecanoic acid (C19:0, 5 nmol) was added as an internal standard to each sample. Total fatty acid methyl esters were prepared using a fatty acid methylation/purification kit (Nacalai Tesque) according to the manufacturer’s instructions. In summary, total fatty acids were extracted using hexane. The esterified fatty acids were saponified and methylated by a sodium hydroxide/methanol solution. Free fatty acids were methylated by a hydrochloric acid/methanol solution. After concentration by evaporation, the methyl ester-derivatized fatty acid was reconstituted with 100 µl of hexane for subsequent analysis. Fatty acids were analyzed using gas chromatography/mass spectrometry (QP2010 Ultra, Shimadzu). The capillary column used for fatty acid separation was SP-2650 (100 m length × 0.25 mm inner diameter × 0.20 µm film thickness, Sigma-Aldrich). The column oven temperature was increased from 140°C to 240°C, and the separated fatty acid methyl ester was detected using mass spectrometry. The standard mixture of methyl ester fatty acids was obtained from Sigma-Aldrich.

### Lipid droplet induction and staining

Oleic acids (Merck) were conjugated with fatty acid-free bovine serum albumin (Fujifilm Wako) as previously described (Ohsaki et al., 2016). For the staining of LDs, Lipi-Blue or Lipi-Deep red (Dojindo Laboratories) was used according to the manufacturer’s instructions.

### Immunofluorescence

Cells were seeded in 35 mm glass-bottom dishes (Matsunami Glass) and incubated under the conditions indicated in each figure legend. Fixation and immunofluorescence staining were performed as previously described (Sakai and Sugasawa, 2014). The following antibodies were used in this study: FANCD2 (NB100-182, Novus Biologicals), phospho-H2AX (05-636, Merck), and PML (sc-966, Santa Cruz Biotechnology). Alexa Fluor secondary antibodies were purchased from Thermo Fisher Scientific. Images were acquired on Olympus FV3000 a confocal laser scanning microscope (Evident) using a UPlanSApo 40× lens with a numerical aperture (NA) of 0.95.

### Immunoblot analysis

Protein extracts were prepared with CSK buffer (10 mM PIPES-NaOH [pH 6.8], 3 mM MgCl_2_, 1 mM EGTA, 300 mM NaCl, 10% glycerol, 0.1% Triton X-100, 50 mM NaF) containing protease inhibitors (0.25 mM phenylmethylsulfonyl fluoride, 1 μg/ml leupeptin, 2 μg/ml aprotinin, 1 μg/ml pepstatin, 50 μg/ml Pefabloc SC). After incubation on ice for 30 min, the cell lysates were centrifuged for 10 min at 20,000 × *g* to obtain soluble extracts. The insoluble fractions were resuspended in the same buffer via sonication and used to detect γH2AX and Lamin B1 in immunoblot analysis. The protein concentration was measured using an XL-Bradford assay kit (Apro Science). Immunoblotting was performed as previously described (Sakai et al., 2020). The following antibodies were used in the present study: FANCD2 (sc-20022, Santa Cruz Biotechnology), FANCI (ab74332, Abcam), phospho-H2AX (A300-081A, Fortis Life Sciences; 05-636, Merck), α-tubulin (T5168, Merck), Lamin B1 (C-20, Santa Cruz Biotechnology), and FLAG (PM020, Medical & Biological Laboratories). Alkaline phosphatase-conjugated secondary antibodies were purchased from Merck.

### Treatment with siRNA

The target sequence for the depletion of FANCD2 has been described previously (Liu et al., 2010), and siRNA was synthesized by Japan Bio Services. The control siRNA (AllStars Negative Control siRNA) was purchased from Qiagen. Cells were transfected with siRNA using the Lipofectamine RNAiMAX reagent (Thermo Fisher Scientific), and the final concentration of siRNA in the culture medium was adjusted to 40 nM.

### Fluorescence recovery after photobleaching

U2OS cells expressing EGFP-FANCD2 were seeded in 35 mm glass-bottom dishes (glass diameter, 27 mm; glass thickness No. 1S; poly-lysine coated; Matsunami Glass) and cultured under the conditions indicated in the legend. Under an FV3000 confocal laser scanning microscope (Evident) equipped with a UPlanSApo 40× lens with a 0.95 NA, EGFP fluorescence was bleached with a 488 nm laser at 100% power. After bleaching, fluorescence images were acquired every 10 s for 5 min.

### Correlative light–electron microscopy

U2OS cells stably expressing EGFP-FANCD2 were cultured on gridded glass coverslips (Matsunami Glass) and treated with 100 µM of OA for 24 h. After labeling with lipid-tox red (Thermo Fisher Scientific) to visualize LDs, the cells were fixed with 4% paraformaldehyde and 0.05% glutaraldehyde in 0.1 M phosphate buffer for 30 min. Fluorescence images were captured by an A1 confocal laser scanning microscope (Nikon) equipped with a GaASP multidetector using a PlanApo 100× lens with a 1.45 NA. Cells were fixed with 2.5% glutaraldehyde in 0.1 M sodium cacodylate buffer (pH 7.4) for 2 h, postfixed with 1% osmium tetroxide and 0.1% potassium ferrocyanide for 1 h, dehydrated, and embedded in epoxy resin. Ultrathin sections were cut and observed using a JEM1400 (JEOL) operated at 80 kV.

### Quantification and statistical analysis

Quantification of the immunoblot analysis was performed using GeneQuant (Cytiva). All signals were normalized by the intensity of α-tubulin or Lamin B1. Statistical analyses were performed as indicated in the figure legends using GraphPad Prism 9 (GraphPad Software).

## Acknowledgements

The authors are grateful to the members of Biosignal Research Center and the Department of Biology, Graduate School of Science, Kobe University, for helpful discussions and encouragement. This work was supported by a Grant-in-Aid to W.S. (KAKENHI Grant Number 17K07286 and 20K06487), and by a Grant-in-Aid to Y.Oh. (KAKENHI Grant Number 21K06733 and 24K02208).

## Author contributions

T. Hotani: formal analysis, investigation, visualization, methodology, and writing—original draft, review, and editing.

M. Goto: formal analysis, investigation, and methodology

Y. Otsuki: formal analysis, investigation, and methodology

S. Matsuda: formal analysis, investigation, and methodology

N. Wada: formal analysis, investigation, and methodology

M. Shinohara: formal analysis, investigation, and methodology

M. Matsuda: resources and supervision

M. Yokoi: resources and supervision

K. Sugasawa: resources and supervision

Y. Ohsaki: conceptualization, supervision, funding acquisition, and writing—original draft, review, and editing

W. Sakai: conceptualization, supervision, funding acquisition, project administration, and writing—original draft, review, and editing

---

**Figure S1.**
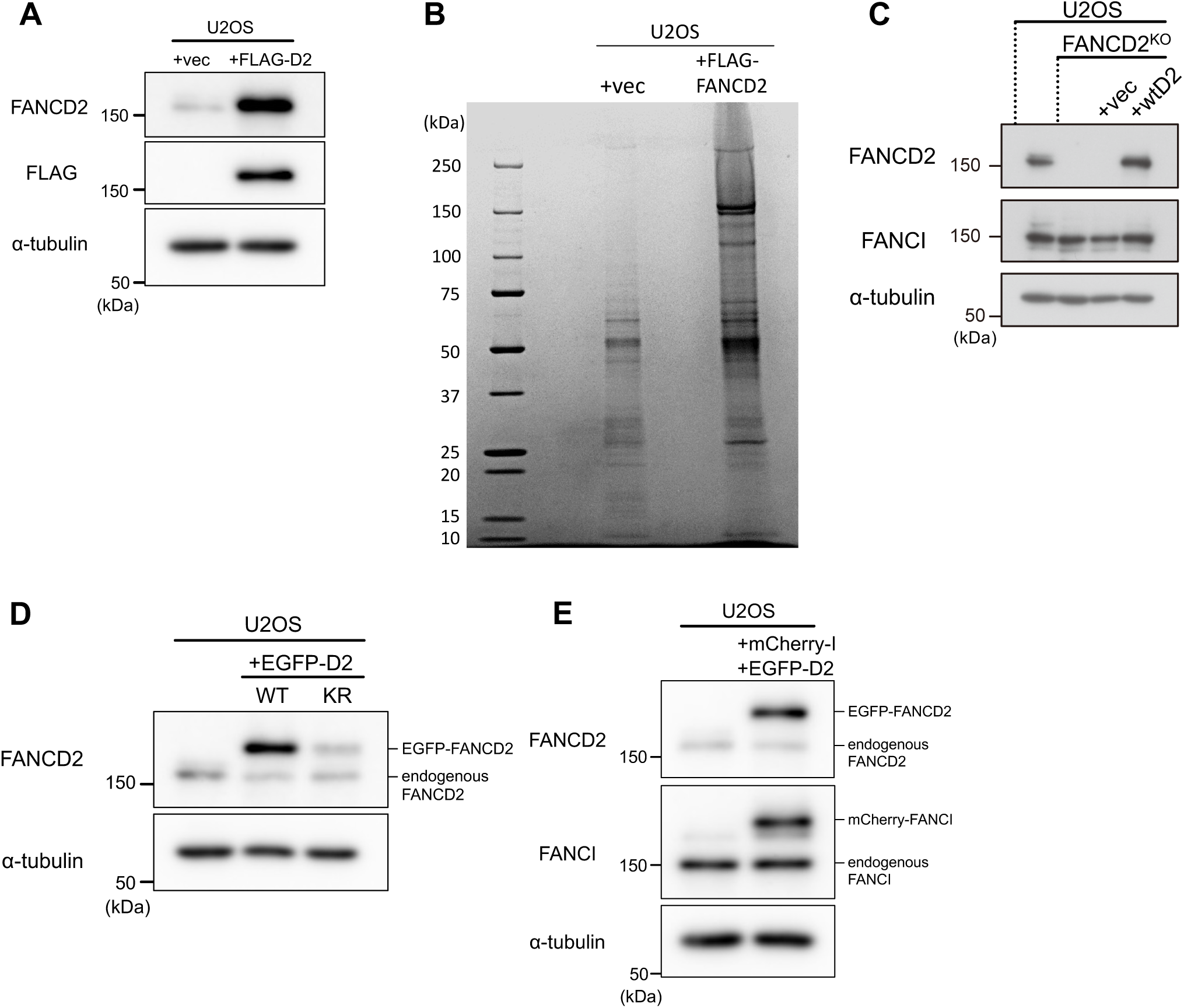
**(A)** Immunoblot analyses confirming the presence of endogenous FANCD2 and ectopically expressed FLAG-FANCD2. **(B)** The FANCD2 complex was immunoprecipitated from U2OS cells expressing FLAG-FANCD2. CBB staining is shown. **(C)** Immunoblot analyses of FANCD2 knockout (FANCD2^KO^) and ectopically expressed FLAG-FANCD2 (wtD2). (**D**) Immunoblot analyses of EGFP-FANCD2 (WT) and mutant EGFP-FANCD2 (KR) in U2OS cells. **(E)** Immunoblot analyses confirming mCherry-FANCI and EGFP-FANCD2 in U2OS cells.

**Figure S2.**
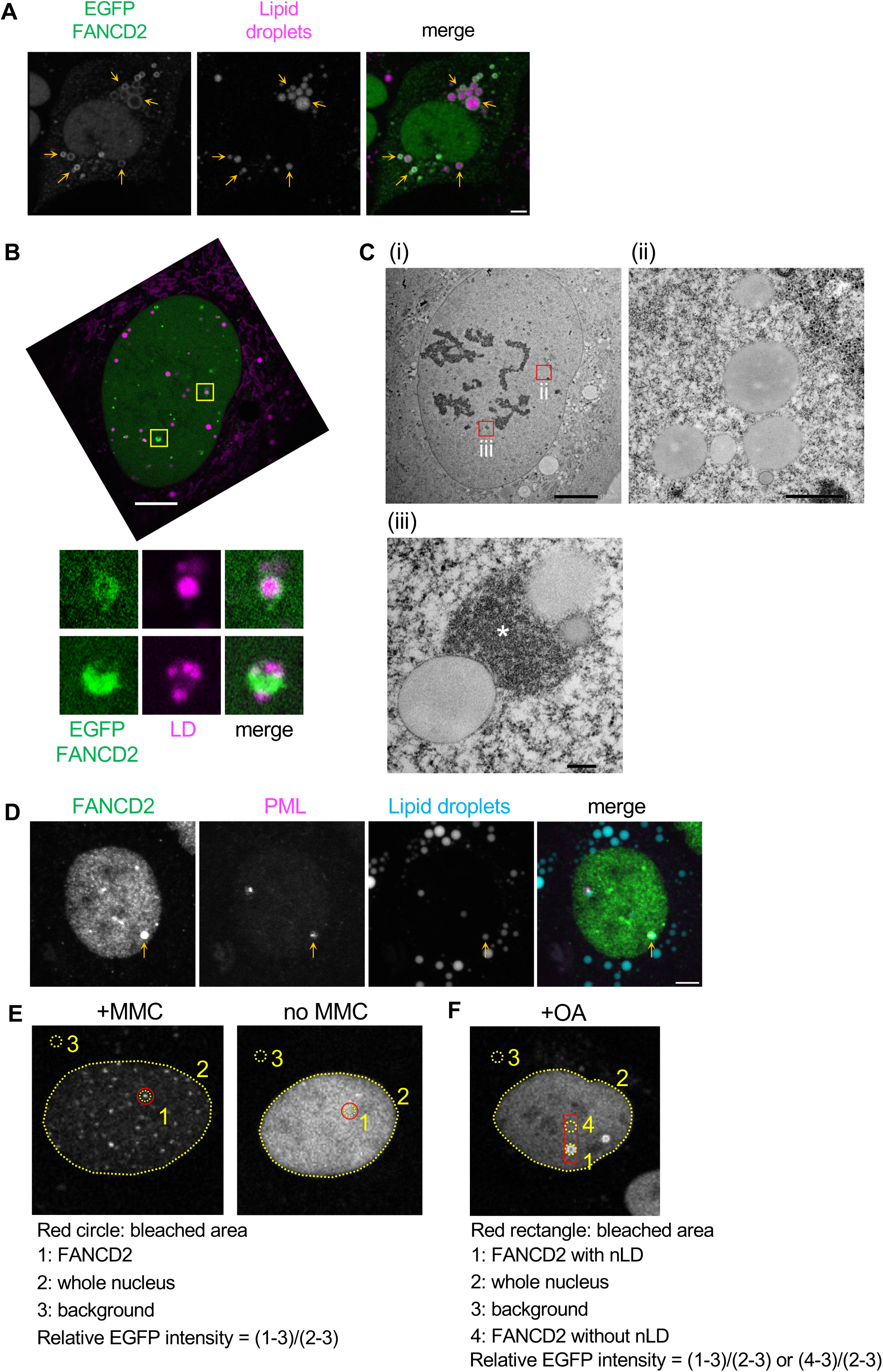
**(A)** Localization of EGFP-FANCD2 in cells cultured with 200 µM of oleic acid (OA) for 2 days. Lipid droplets (LDs) were stained with Lipi-Blue. Scale bar, 5 µm. (**B**, **C**) Correlative light–electron microscopy. U2OS cells stably expressing EGFP-FANCD2 were treated with 100 µM of OA for 24 h. (**B**) Cells were weakly fixed, and fluorescence images were captured first. Some nuclear LDs (nLDs; magenta) were surrounded by FANCD2 (green). Highly magnified images (yellow square) are shown below. The upper and lower panels correspond with C (ii) and (iii), respectively. Scale bar, 10 µm. (**C**) (i) Electron microscopy images of the cell shown in (**B**). (ii) and (iii) Highly magnified images (red squares) are shown. Scale bars, 10 µm (i), 1 µm (ii), and 200 nm (iii). The asterisk indicates the condensed chromatin area. (**D**) Localization of endogenous FANCD2 and PML in U2OS cells cultured with 200 µM of OA for 2 days. Orange arrows indicate an nLD with FANCD2 and PML. Representative images are shown. Scale bar, 5 µm. **(E, F)** The relative EGFP-FANCD2 intensity in the FRAP analysis was calculated using the indicated formulas. The red circles and rectangle indicate the bleached areas. The numbers in panels E and F indicate the following: (**E**) 1, FANCD2; 2, whole nucleus; 3, background and (**F**) 1, FANCD2 with LD; 2, whole nucleus; 3, background; 4, FANCD2 without LD.

**Figure S3.**
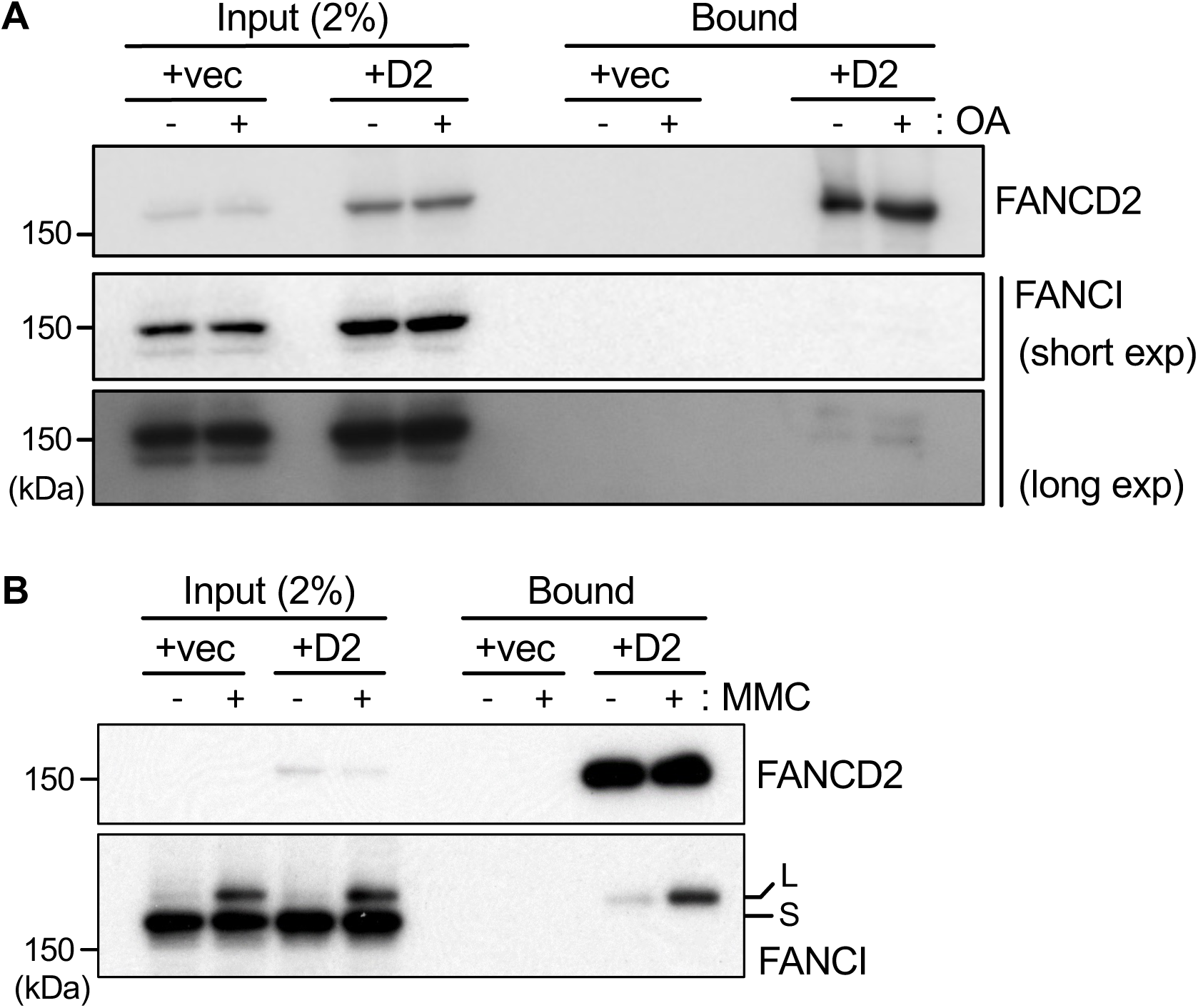
(**A, B**) Co-immunoprecipitation assays were performed using U2OS cells stably expressing FLAG-FANCD2. Total cell extracts were prepared from cells treated with 200 µM oleic acid (OA) for 2 days or 1 µM mitomycin C (MMC) overnight and subjected to immunoprecipitation with anti-FLAG beads. (**B**) L and S indicate the monoubiquitinated and unmodified forms of FANCI, respectively.

**Videos 1 and 2**

EGFP-FANCD2-expressed U2OS was treated with 200 µM oleic acid (OA) for 24 h. One frame was captured every 5 min. Scale bar, 5 µm. Selected frames are shown in Fig 2AB.

